# SMC progressively aligns chromosomal arms in *Caulobacter crescentus* but is antagonized by convergent transcription

**DOI:** 10.1101/125344

**Authors:** Ngat T. Tran, Michael T. Laub, Tung B. K. Le

## Abstract

The Structural Maintenance of Chromosomes (SMC) complex plays an important role in chromosome organization and segregation in most living organisms. In *Caulobacter crescentus*, SMC is required to align the left and the right arms of the chromosome that run in parallel down the long axis of the cell. However, the mechanism of SMC-mediated alignment of chromosomal arms remains elusive. Here, using a genome-wide chromosome conformation capture assay, chromatin immunoprecipitation with deep sequencing, and microscopy of single cells, we show that *Caulobacter* SMC is recruited to the centromeric *parS* site and that SMC-mediated arm alignment depends on the chromosome partitioning protein ParB. We provide evidence that SMC likely tethers the *parS-*proximal regions of the chromosomal arms together, promoting arm alignment. Strikingly, the co-orientation of SMC translocation and the transcription of highly-expressed genes is crucial for the alignment of *parS-*proximal regions of the chromosomal arms. Highly-transcribed genes near *parS* that are oriented against SMC translocation disrupt arm alignment suggesting that head-on transcription interferes with SMC translocation. Our results demonstrate a tight interdependence of bacterial chromosome organization and global patterns of transcription.

## INTRODUCTION

The chromosomes of all organisms must be compacted nearly three orders of magnitude to fit within the limited volume of a cell. However, DNA cannot be haphazardly packed, and instead it must be organized in a way that is compatible with numerous cellular processes that share the same DNA template, including transcription, DNA replication, and chromosome segregation. This is particularly challenging in bacteria since these chromosome-based transactions happen concomitantly rather than being separated temporally, as in eukaryotes. Application of microscopy-based analyses of fluorescently-labeled DNA loci along with genome-wide chromosome conformation capture assays (Hi-C) have revealed a well-defined *in vivo* three-dimensional organization of bacterial chromosomes (Badrinarayanan et al., 2015a; Le et al., 2013; Umbarger et al., 2011; Viollier et al., 2004). Hi-C provides quantitative information about the spatial proximity of DNA loci *in vivo* by measuring the frequencies of crosslinking between different regions of the chromosome (Lieberman-Aiden et al., 2009). The first application of Hi-C to bacteria examined the *Caulobacter crescentus* chromosome (Le et al., 2013). Hi-C analysis of *Caulobacter* confirmed the global pattern of chromosome organization: in cells with a single chromosome, the origin of replication (*ori*) is at one cell pole, the terminus (*ter*) is near the opposite pole, and the two chromosomal arms are well-aligned, running in parallel down the long axis of the cell (Le et al., 2013; Viollier et al., 2004) (Fig. 1A). We discovered that a Structural Maintenance of Chromosomes protein (SMC) is crucial for the alignment of the left and right arm of the chromosome in *Caulobacter* (Le et al., 2013), but how SMC achieves this alignment remains poorly understood.

**Figure 1:**
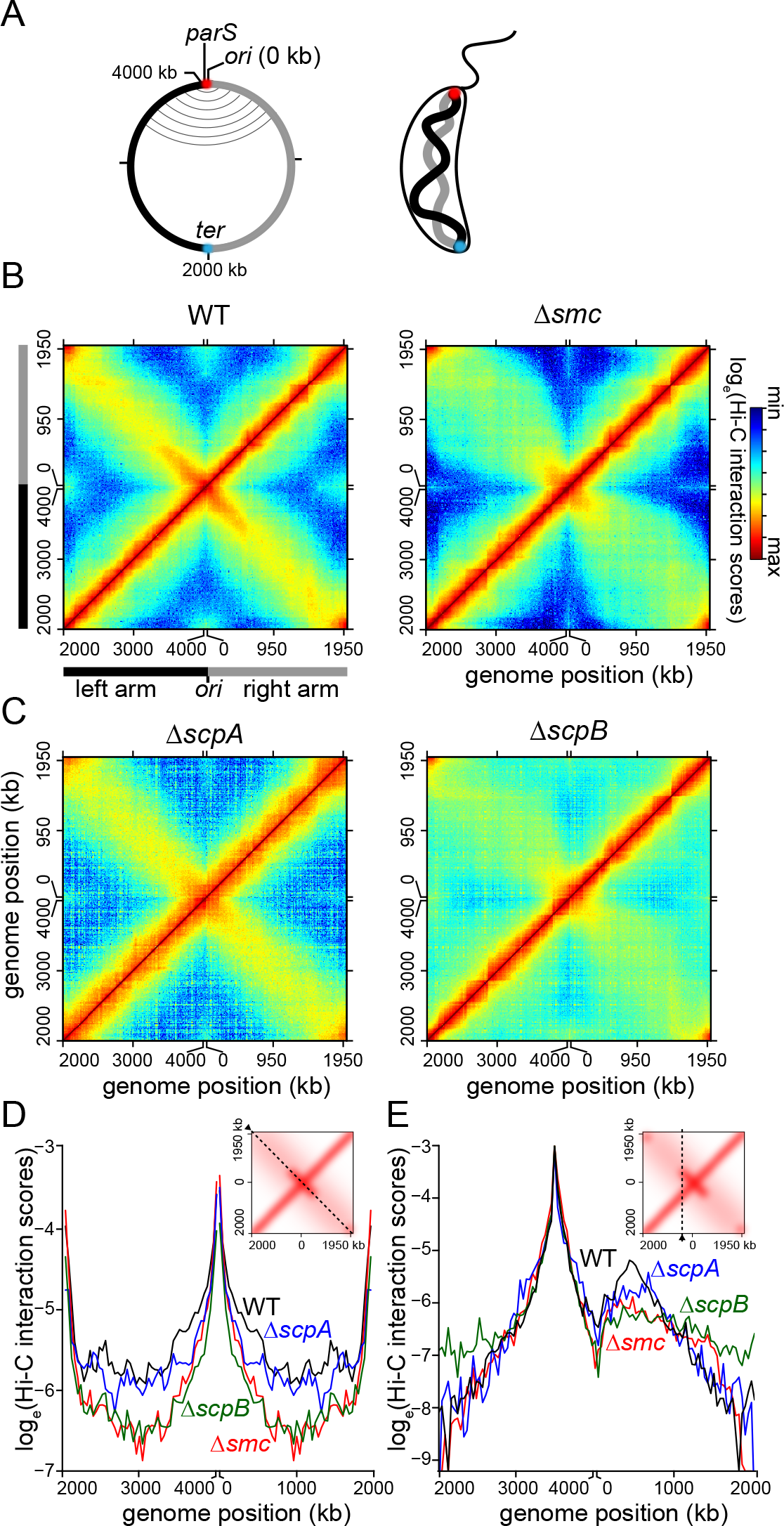
The SMC-ScpA-ScpB complex promotes the alignment of chromosomal arms. **(A)** A simplified genomic map of *Caulobacter* showing the origin of replication (*ori*), the *parS* site and the terminus (*ter*), together with left (black) and the right (grey) chromosomal arms. On the genomic map, aligned DNA regions are presented schematically as grey curved lines connecting the two chromosomal arms. Spatially, *ori* (red) and *ter* (cyan) are at opposite poles and the two arms running in parallel down the long axis of the cell. **(B)** Normalized Hi-C contact maps showing the natural logarithm of DNA-DNA contacts for pairs of 10 kb-bins across the genome of WT and *Δsmc* cells (Le et al., 2013). **(C)** Normalized Hi-C contact maps for *ΔscpA* and *ΔscpB* cells. **(D)** Hi-C interaction scores along the secondary diagonal (black dashed line in the inset) for contact maps of WT, *Δsmc, ΔscpA* and *ΔscpB* cells. Bins near *ori* or near *ter* are dominated by intra-arm instead of inter-arm DNA-DNA interactions due to the circular nature of the chromosome. **(E)** Hi-C interaction scores along the vertical line (black dashed line in the inset) for contact maps of WT, *Δsmc, ΔscpA* and *ΔscpB* cells.

SMC proteins are widely conserved from bacteria to humans. In eukaryotes, SMC1 and SMC3, together with accessory proteins, form a cohesin complex that holds sister chromatids together until after they achieve bipolar attachment to the mitotic spindle. The related condensin complex, comprised of SMC2, SMC4, and interacting partners, promotes the compaction of individual chromosomes during mitosis. In most bacteria, there is a single SMC composed of an ATPase “head” domain, a dimerization “hinge” domain, and an extended antiparallel coiled-coil region in the middle (reviewed in Nolivos and Sherratt, 2014). Two SMC monomers dimerize, and together with the bacteria-specific proteins ScpA and ScpB, form a tripartite ring that can bring distal DNA segments close together to help organize bacterial chromosomes (Bürmann et al., 2013; Mascarenhas et al., 2002). The topological entrapment of DNA by a ring-shaped SMC complex has been shown for cohesin and condensin, and for *Bacillus subtillis* SMC (Cuylen et al., 2011; Ivanov and Nasmyth, 2005; Wilhelm et al., 2015). It is likely that topological entrapment is a general feature of all SMC complexes.

How SMC gets loaded onto DNA and topologically entraps DNA is generally well-studied in eukaryotes but not yet completely understood (reviewed in Uhlmann, 2016). In bacteria, SMC loading, translocation and DNA entrapment is less well known, but likely involves the ParA-ParB-*parS* system (Gruber and Errington, 2009; Lin and Grossman, 1998; Minnen et al., 2011). ParB is a DNA-binding protein that nucleates on a centromere-like *parS* sequence (Mohl and Gober, 1997) and then spreads non-specifically along the DNA, likely forming a large nucleoprotein complex (Breier and Grossman, 2007; Graham et al., 2014). ParA, a Walker-box ATPase, interacts with ParB and is required for the segregation of replicated chromosomes to daughter cells (Figge et al., 2003). In *B. subtilis*, ParB loads SMC onto the chromosome mainly at the *ori*-proximal *parS* sites (Gruber and Errington, 2009; Marbouty et al., 2015; Wang et al., 2015). Loaded SMC then translocates from *parS* to distal parts of the chromosome in an ATP hydrolysis-dependent manner (Minnen et al., 2016; Wang et al., 2017). This action is thought to individualize the origins of replicated chromosomes, thereby helping to segregate replicated chromosomes. It has been proposed that *B. subtilis* SMC loaded at *parS* translocates the full length of the chromosome to *ter* to promote chromosome arm alignment (Gruber and Errington, 2009; Minnen et al., 2016; Wang et al., 2017).

However, ChIP-seq studies indicate that *B. subtilis* SMC is most enriched near *ori*, so whether SMC directly promotes arm alignment uniformly across the genome is unclear (Minnen et al., 2016; Wang et al., 2017).

*Caulobacter* harbours both a canonical SMC-ScpA-ScpB complex as well as a ParA-ParB-*parS* system. The ParA-ParB-*parS* complex is essential (Mohl and Gober, 1997). In contrast, *Caulobacter* SMC is not required for survival in laboratory conditions (Le et al., 2013). *Caulobacter* cells lacking SMC grow slightly slower but are not temperature sensitive nor prone to accumulating suppressor mutations as originally suggested (Jensen and Shapiro, 1999). Nevertheless, ectopic overexpression of an ATPase-defective *smc* mutant shows a severe defect in sister chromosome segregation in this bacterium, consistent with *Caulobacter* SMC playing a role in chromosome segregation (Schwartz and Shapiro, 2011). How SMC influences other cellular processes like transcription and is, in turn, influenced by these processes, is not well understood. In yeast, cohesin is pushed along the chromosome in the same direction as transcription, without dissociating (Ocampo-Hafalla et al., 2016). Similarly, in budding yeast, RNA polymerase can drive the short-range relocation of condensin (D’Ambrosio et al., 2008; Johzuka and Horiuchi, 2007). Whether bacterial RNA polymerase affects SMC has not been systematically investigated. Notably, in almost all bacteria, most genes, especially highly-expressed genes, are transcribed from *ori* to *ter* (reviewed in Rocha, 2008). This co-orientation of genes could help push SMC toward *ter*; alternatively, or in addition, genes transcribed in a head-on orientation could antagonize the translocation of SMC.

Here, we use Hi-C and ChIP-seq, together with microscopy-based analysis of single cells to elucidate the role and the mechanism of SMC in the global organization of the *Caulobacter* chromosome. We provide evidence that (i) SMC is required for the progressive alignment of the two chromosomal arms, proceeding in the *ori-ter* direction; (ii) *Caulobacter* SMC is loaded onto the chromosome at the *parS* site and ParB is essential for the SMC-mediated arm alignment; (iii) *Caulobacter* SMC most likely functions as a tether to actively cohese ~600 kb *parS-*proximal regions of the chromosomal arms together; and (iv) head-on transcription can profoundly disrupt the alignment of chromosomal arms, likely by interfering with SMC translocation from *parS*. Altogether, our results demonstrate a tight interdependence of bacterial chromosome organization and global patterns of transcription.

## RESULTS

### The SMC-ScpA-ScpB complex is required for the alignment of chromosomal arms

*Caulobacter* cells lacking SMC show a dramatic reduction in inter-arm DNA-DNA interactions (Le et al., 2013) (Fig. 1B). To test whether the *Caulobacter* ScpA and ScpB homologs are also required for inter-arm interactions we generated Hi-C contact maps for a homogenous G1-phase populations of Δ*scpA* and Δ*scpB* cells (Fig. 1B-C). We divided the *Caulobacter* genome into 10-kb bins and assigned to corresponding bins the interaction frequencies of informative ligation products. Interaction frequencies were visualized as a matrix with each matrix element, *m_ij_*, indicating the natural logarithm of the relative interaction frequency of DNA loci in bin *i* with those in bin *j*. To emphasize the *ori*-proximal region, we oriented the Hi-C contact maps such that the *ori* (0 kb or 4043 kb) is at the center of the x- and y-axis, and the left and the right chromosomal arm are on either side (Fig. 1B-C).

On the contact map of wild-type (WT) *Caulobacter*, the primary and high-interaction diagonal represents interactions between DNA on the same arm of the chromosome, *i.e*. intra-arm contacts, while the less prominent secondary diagonal represents DNA-DNA interactions between opposite arms, *i.e*. inter-arm contacts (Fig. 1B). The diagonal pattern of inter-arm contacts on a Hi-C map indicates that each locus on one chromosomal arm interacts with a locus roughly equidistant from the *ori* on the opposite arm, reflecting a global alignment of the chromosomal arms. Consistent with our previous studies (Le et al., 2013), the inter-arm interactions are significantly reduced in a Δ*smc* strain. The Hi-C map for Δ*scpB* revealed a similar decrease in inter-arm interactions, with no strong or obvious change in intra-arm interactions (Fig. 1C-E). The Hi-C map for Δ*scpA* also exhibited a decrease in inter-arm interactions, though not nearly as significant as for Δ*smc* and Δ*scpB* (Fig. 1C-E). These data are consistent with ScpA and ScpB forming a complex with SMC that promotes the co-linearity of chromosomal arms in *Caulobacter*.

### ParB induces a progressive alignment of chromosomal arms from ori to ter

In Gram-positive bacteria such as *B. subtilis* and *Streptococcus pneumoniae*, SMC is loaded onto the chromosome by ParB at *ori*-proximal *parS* sites (Gruber and Errington, 2009; Minnen et al., 2011). To test whether this mechanism is conserved in *Caulobacter*, a Gram-negative bacterium, we used a strain where ParB, which is essential for viability, can be depleted. The promoter of *parB* at its native chromosomal locus was replaced with a xylose-inducible promoter, P_*xyl*_. Cells grown to exponential phase in the presence of xylose were then washed free of xylose and incubated for five hours in a rich medium supplemented with glucose to inhibit P_*xyl*_ activity. Immunoblot analysis with an α-ParB antibody indicated ~2.5 fold decrease in ParB concentration after the five hours in glucose (T = 0 min, Fig. S1A). The contact map of ParB-depleted cells exhibited a clear reduction in inter-arm contacts, similar to Δ*smc* cells, indicating a role of ParB in maintaining chromosomal arm alignment, possibly by loading SMC onto DNA (Fig. 2A-B).

**Figure 2:**
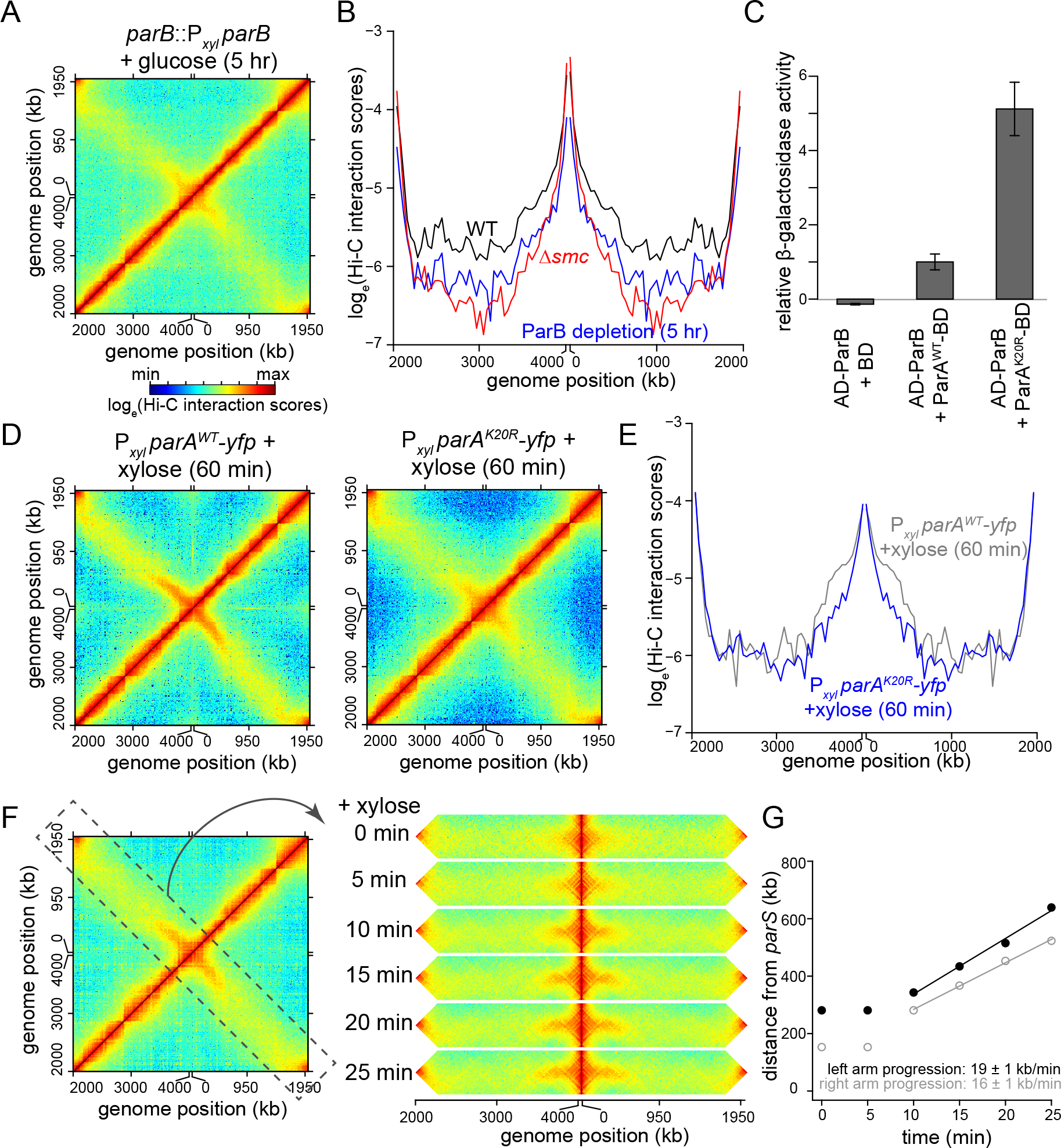
ParB is required for the progressive alignment of chromosomal arms by SMC. **(A)** Normalized Hi-C maps for *parB::*P*xyl parB* cells 5 hrs after starting the depletion experiment. **(B)** Hi-C interaction scores along the secondary diagonal for contact maps of WT (black), *Δsmc* (red), and *parB::*P_*xyl*_*parB*cells (blue) 5 hrs after starting the depletion experiment. **(C)** Yeast two hybrid assay to compare ParB-ParA^WT^ interaction to that of ParB-ParA^K20R^. The β-galactosidase activity was assayed for each strain and is presented relative to the value obtained for the ParB-ParA^WT^ interaction. Error bars represent standard deviation from four biological replicates. **(D)** Normalized Hi-C maps for cells over-expressing *parA^WT^-yfp* and *parA^K20R^-yfp* after adding xylose for 1 hr. **(E)** Hi-C interaction scores along the secondary diagonal for contact maps of cells over-expressing *parA^WT^-yfp* (grey), and *parA^K20R^-yfp* (blue) after adding xylose for 1 hr. **(F)** A time-resolved Hi-C for cells that are replenishing of ParB. Time after adding back xylose was indicated next to each Hi-C strip. For presentation purposes, the secondary diagonal (black dashed box) was rotated and laid out horizontally. **(G)** Analysis of the progression of chromosomal arm alignment. The extent of DNA alignment at each time point after adding back xylose was plotted for each chromosomal arm. The black and grey lines are linear best fit lines for data from time point 10 min to 25 min.

As with *B. subtilis* SMC (Gruber and Errington, 2009; Wang et al., 2015), the *Caulobacter* SMC complex may be recruited and loaded onto DNA via a direct interaction with ParB. If so, we reasoned that overexpressing a strong ParB-interacting protein might prevent interactions with SMC and, in turn, disrupt alignment of the chromosomal arms. To test this hypothesis, we performed Hi-C on cells overexpressing a YFP-tagged variant of ParA harboring a K20R substitution. ParA(K20R) is defective in ATP binding but binds ParB more tightly than ParA (WT), at least in a yeast two-hybrid assay (Fig. 2C and Shebelut et al., 2010). The Hi-C contact map for a strain overexpressing ParA(K20R) showed a modest, but significant decrease in chromosomal arm alignment compared to a control strain overexpressing ParA (WT) (p <10^-12^, paired Student’s t-test, Fig. 2D-E, Fig. S1B-C). This result reinforces our conclusion that ParB is required for the SMC-mediated alignment of chromosomal arms in *Caulobacter*.

To investigate the directionality and dynamics of chromosome arm alignment, we again depleted ParB by growing cells in rich medium with glucose for 5 hours, and then added back xylose to induce ParB *de novo*. Samples were taken 0, 5, 10, 15, 25 and 30 minutes after xylose addition for Hi-C analysis. Immunoblot analysis with α-ParB antibody showed gradual accumulation of new ParB after adding back xylose (Fig. S1A). At the 0 and 5 minute time points, we observed very little inter-arm interaction, as above (Fig. 2F, Fig. S1D-E). However, over time, the inter-arm interactions increased, beginning close to *ori* and *parS* and then extending toward *ter* (Fig. 2F-G, Fig. S1D-E). The two arms aligned directionally at a rate of ~19 kb per minute and ~16 kb per minute for the left and right arm, respectively (Fig. 2F-G). These data are consistent with a model in which SMC is loaded by ParB, likely at *ori/parS*, and then translocates down the arms toward *ter*, driving their alignment. Nevertheless, we cannot formally exclude the possibility that ParB might have additional, SMC-independent roles in chromosomal arm alignment.

The *parS* site is ~8 kb from *ori* in *Caulobacter* (Toro et al., 2008). To directly test the model that SMC is loaded onto DNA at *parS* sites, we inserted a 260-bp DNA fragment containing a *parS* site either at +1800 kb or +2242 kb from *ori* (Fig. S2A). We verified that this ectopic *parS* site was sufficient to recruit ParB, using ChIP-seq analysis of ParB, which binds to the native *parS* site (Fig. S2B). We then performed Hi-C on the strain harboring the additional *parS* site and observed a new secondary diagonal, emanating from the approximate position of the ectopic *parS* site (Fig. S2C). The extent of alignment from this ectopic site was less than that associated with the original *parS*, an issue examined in depth below. Taken together, our results support a model in which ParB loads SMC at *parS* sites, leading to the subsequent progressive alignment of flanking DNA.

### SMC promotes DNA alignment most effectively for parS-proximal genomic regions

After being loaded at *parS* sites, SMC likely translocates down the chromosomal arms to drive their alignment. However, the extent of inter-arm interactions was not uniform across the entire chromosome indicating that each loaded SMC complex may not travel the entire length of the chromosome. In fact, we noted that in WT cells, inter-arm interactions reduced gradually away from *parS*, leveling out after ~600 kb in either direction (Fig. 1D). These observations suggested that SMC is not uniformly distributed across the chromosome and instead may be enriched in the regions showing highest inter-arm interactions. To test this hypothesis, we first used ChIP-seq to map the genome-wide distribution of epitope tagged-SMC. We fused the SMC-encoding gene to a FLAG tag at the N-terminus and placed this allele downstream of P_*xyl*_ on a medium-copy-number plasmid. We then performed Hi-C on *Δsmc* cells that produced this FLAG-tagged SMC via leaky expression from P_*xyl*_ (Fig. S3A). Chromosomal arm alignment was comparable to the WT level (Fig. S3B-C), indicating that FLAG-SMC is functional. Overproducing FLAG-SMC in the *Δsmc* background by adding xylose did not extend arm alignment beyond the WT level (Fig. S3B-C). We performed α-FLAG ChIP-seq with the *Δsmc* P_*xyl*_-*flag-smc* strain. As a negative control, we performed α-FLAG ChIP-seq on WT *Caulobacter, i.e*. cells with untagged SMC (Fig. S3D). Although only a small percentage of ChIP DNA is enriched by FLAG-SMC, we observed a clear enrichment of SMC-bound DNA near the *parS* site (Fig. 3A), consistent with SMC loading at *parS*. SMC decreases in enrichment away from *parS* but is enhanced close to the *ter* area (Fig. 3A and the Discussion). We also noted the enrichment of SMC at highly transcribed genes (Fig. 3A-B), most likely as an artefact of non-specific immunoprecipitation (Teytelman et al., 2013).

**Figure 3:**
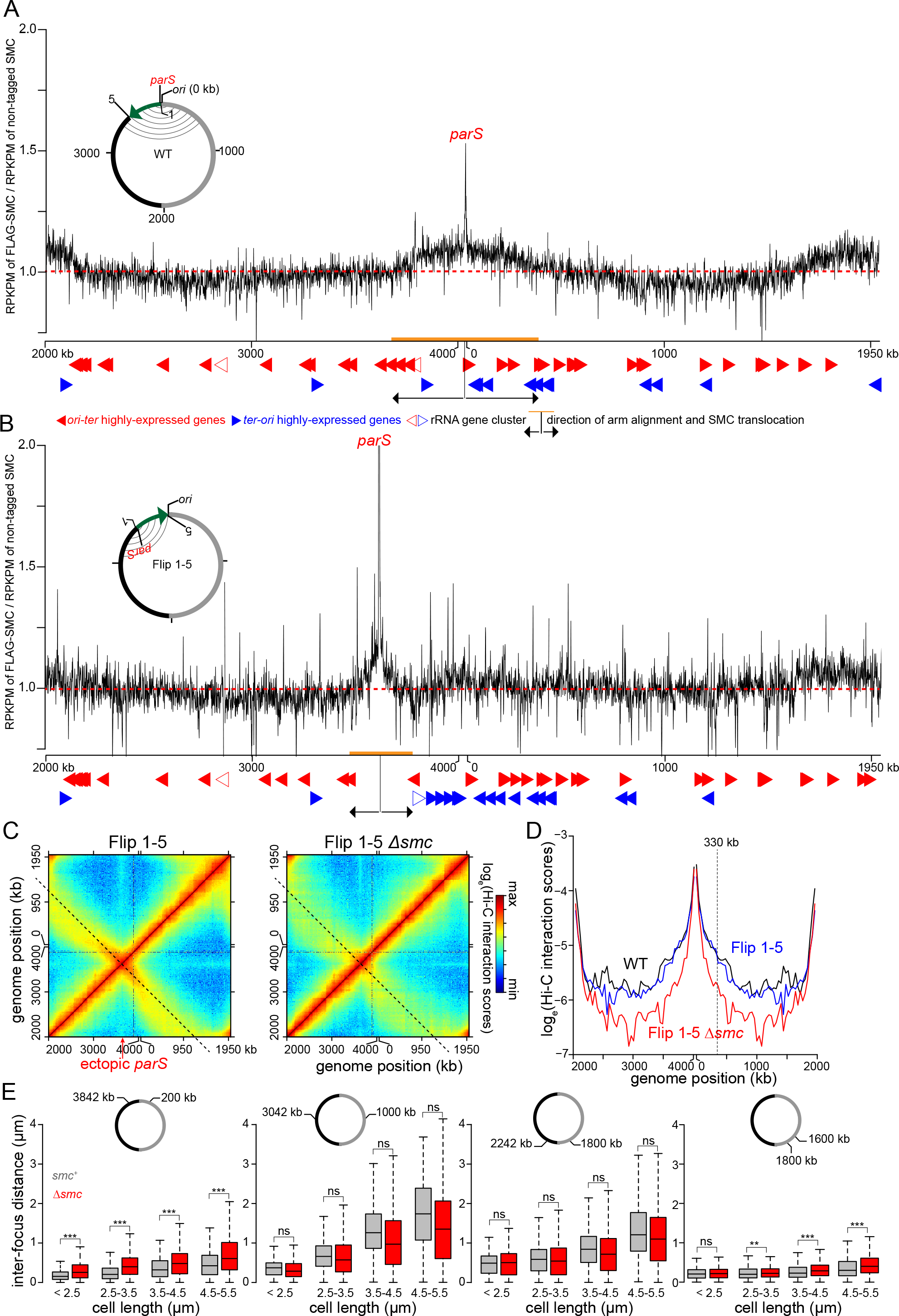
Fig. 3. SMC is enriched at the *parS* site and promotes DNA alignment most effectively for *parS*-proximal regions. **(A)**The distribution of FLAG-tagged SMC on WT *Caulobacter* chromosome. ChIP-seq signals were reported as the number of reads within every 1 kb bin along the genome (RPKPM) in the ChIP fraction of FLAG-tagged SMC divide by untagged SMC. The dashed red line shows y-axis value at 1. Below the ChIP-seq profile are the position of highly-expressed genes that transcribe in the *ori-ter* (solid red arrows) or *ter-ori* (solid blue arrows) direction. The positions of ribosomal RNA gene cluster are indicated with open red or blue arrows. The direction and extent of SMC translocation from *parS* site were shown as black arrows and orange bar, respectively. A schematic genomic map of *Caulobacter* showing the position of *parS* (red) and *ori* are presented in the inset. The inverted DNA segment (green arrow) is indicated together with the end points of the inversion (1 and 5). On the genomic map, aligned DNA regions, as observed by Hi-C, are presented schematically as grey curved lines connecting the two chromosomal arm. **(B)** The distribution of FLAG-tagged SMC on the chromosome of Flip 1-5 *Caulobacter*. **(C)** Normalized Hi-C maps for the Flip 1-5 and Flip 1-5 *Δsmc* cells. **(D)** Hi-C interaction scores along the secondary diagonal for contact maps of WT (black), Flip 1-5 (blue), and Flip 1-5 *Δsmc* cells (red). The Hi-C interaction scores along the secondary diagonal of Flip 1-5 and Flip 1-5 *Δsmc* cells (black dashed lines in panel **C**) was shifted to the same position as that of WT to enable comparison between strains. The vertical black dashed line at ~330 kb away from *ori* shows the position where Hi-C interaction scores along the secondary diagonal start to reduce in Flip 1-5 strain in comparison to WT. **(E)** Inter-focus distances expand differentially in elongated *Caulobacter* cells, depending on their genomic locations. Pairs of DNA loci were labeled with YFP-ParB^pMT1^/*parS*^pMT1^ and mCherry-ParB^P1^/*parS*^P1^ near *ori* (+200 kb and +3842 kb), at the middle of each arm (+1000 kb and +3042 kb), near *ter* (+1800 kb and +2242 kb), or on the same arm (+1600 kb and +1800 kb). Boxplots show the distribution of inter-focus distances for cells of different sizes with SMC (grey) or without (red) SMC. Asterisks indicate statistical significance (*** P-value < 0.001; ns: not significant, one-tailed Student’s t-test. Null hypothesis: inter-focus distance in *Δsmc* is greater than in WT cells).

Notably, FLAG-tagged SMC was enriched above background in a region overlapping *parS* that extended from approximately +3680 kb to +345 kb, the same approximate region that shows the most extensive inter-arm alignment by Hi-C (Pearson’s correlation coefficient = 0.75, p < 10^-12^, Fig. 3A, Fig. S4A). We could not detect significant enrichment of FLAG-tagged SMC beyond this *parS*-proximal region, indicating that SMC is either not appreciably bound to *parS-*distal regions of the chromosome or its association drops below our limit of detection with ChIP-seq. In either case, we conclude that (i) SMC is not uniformly distributed across the genome and (ii) the enrichment of SMC correlates with the extent of chromosome arm alignment at the *ori*-proximal region.

**Figure 4:**
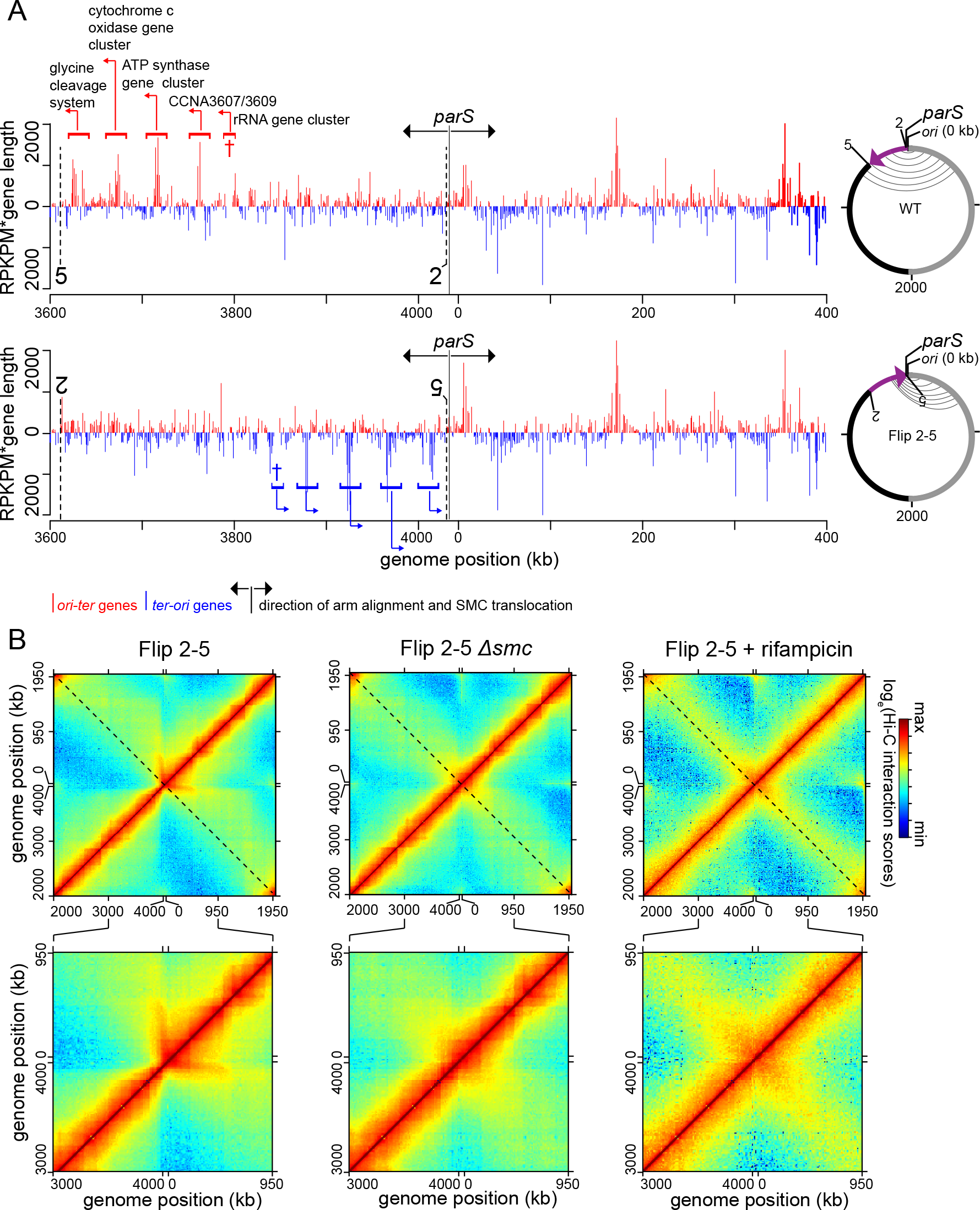
Genomic context and transcription influence the SMC-mediated alignment of chromosomal arms. **(A)** The abundance of RNA polymerases on genes that transcribe in the *ori-ter* direction (red) or in the *ter-ori* direction (blue) in WT and Flip 2-5 cells for the DNA segment between +3600 kb and +400 kb. The position of *parS* and the direction of SMC translocation are indicated with black arrows. For the whole genome plot, see Fig. S6. ChIP-seq using α-FLAG antibody was performed on cells expressing *rpoC-flag* from its native locus in WT background (upper panel) or in Flip 2-5 background (lower panel). The abundance of RNA polymerases was represented as RPKPM*gene length for each gene and plotted against the genomic location of that gene. Due to short sequencing reads and the high similarity between the two rRNA clusters, it is not reliable to estimate the RNA polymerase density within each rRNA cluster. Therefore, enrichment data for rRNA gene clusters are not shown. Nevertheless, we indicate the genomic position of a highly-expressed rRNA cluster on *Caulobacter* genome with a dagger (†) symbol. Vertical black dashed lines with numbering 2 and 5 indicate the inversion end points. **(B)** Normalized Hi-C contact maps for Flip 2-5, Flip 2-5 *Δsmc* cells, and Flip 2-5 cells treated with rifampicin. A 1000 kb region surrounding *parS/ori* were also zoomed in and presented below each whole-genome Hi-C contact map.

To further test the relationship of SMC enrichment and the alignment of flanking DNA, we generated a strain bearing a relatively large inversion (involving genomic DNA normally between +3611 and +4038 kb) such that *parS* is relocated ~427 kb away from *ori*; this strain is referred to as the Flip 1-5 strain (Fig. 3B). ChIP-seq of FLAG-tagged SMC in the Flip 1-5 inversion strain showed an enrichment of DNA surrounding the relocated *parS* site at a peak level comparable to the *ori*-proximal *parS* (Fig. 3A-B), further supporting the conclusion that SMC is loaded at *parS*, and that SMC enrichment near *parS* is independent of *ori*.

The Hi-C contact map of the Flip 1-5 inversion strain also showed a secondary diagonal (Fig. 3C and Fig. S5A). However, the starting point of the flanking DNA alignment was shifted and coincided with the genomic position of the relocated *parS* (Fig. 3C and Fig. S5A). This ectopic arm alignment was reduced dramatically in the absence of *smc* (Fig. 3C-D), indicating that arm alignment in the Flip 1-5 strain still depends on SMC. For the Flip 1-5 strain, the inter-arm alignment was again strongest in a limited region around *parS*, extending approximately 330-420 kb in each direction (Fig. 3D, Fig. S5B). The extent of arm alignment in the Flip 1-5 strain was reduced compared to WT cells (~600 kb), consistent with the reduced region showing SMC enrichment by ChIP-seq (Fig. 3B, Fig. S4B).

**Figure 5:**
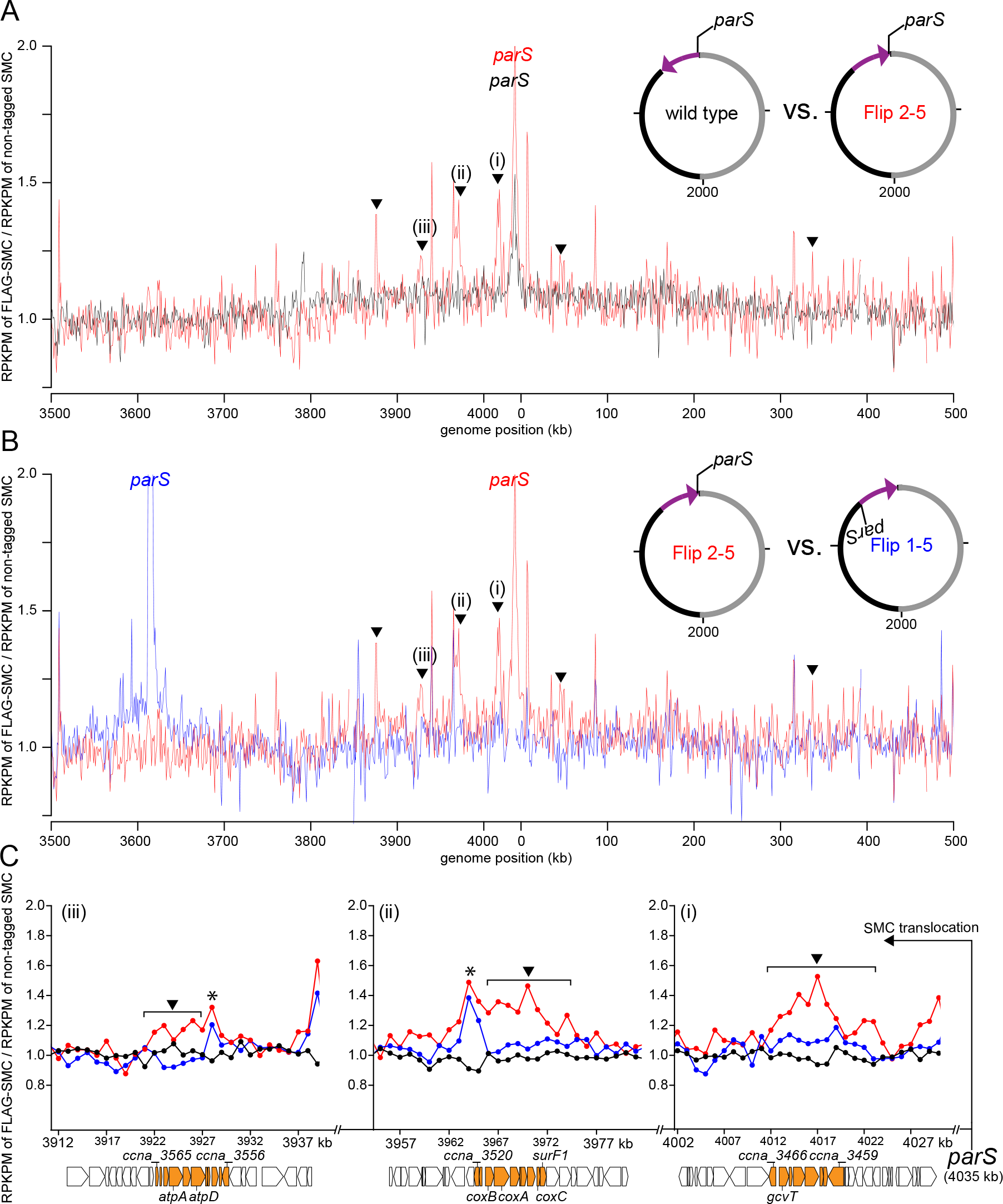
Head-on transcription alters the distribution of SMC on the chromosome. **(A)** The distribution of FLAG-tagged SMC on WT *Caulobacter* chromosome (black) and on Flip 2-5 chromosome (red). Only DNA segment between +3500 kb and +500 kb was shown. For profiles of the whole genome, see Fig. S4. **(B)** The distribution of FLAG-tagged SMC on Flip 1-5 (blue) and Flip 2-5 chromosome (red). Black triangles indicate new peaks in the ChIP-seq profile of Flip 2-5 but not in the profiles of WT or Flip 1-5 strain. The ChIP-seq profile of FLAG-SMC in Flip 1-5 strain was slightly shifted to align to that of Flip 2-5 strain since the inverted DNA segment in Flip 1-5 is larger than in Flip 2-5 by 8 kb to encompass the native *parS* region (See Fig. 4A-B and Fig. 4A). **(C)** The distribution of FLAG-tagged SMC at the glycine cleavage system gene cluster (panel i, highlighted in orange), the cytochrome c oxidase gene cluster (panel ii, highlighted in orange) and the ATP synthase gene cluster (panel iii, highlighted in orange). The genomic position of *parS* and the direction of SMC translocation are shown with a black arrow. The genomic positions on the x-axis and the gene direction are those of Flip 2-5 strain. We inverted *in silico* the ChIP-seq profile and gene orientation of WT strain to enable comparison of superimposed ChIP-seq profiles. Black triangles indicate new peaks in the ChIP-seq profile of Flip 2-5 but not in the profiles of WT or Flip 1-5 strain. Black asterisks (*) indicate non-specific peaks that often associate with “hyper-ChIPable” regions at highly-transcribed genes.

Collectively, the data presented thus far suggest that SMC loaded at *parS* may only translocate a limited distance down the chromosome. If so, we predicted that if the chromosome expanded, as occurs in elongating, G1-arrested cells, then *parS*-proximal regions would (i) remain better aligned than other chromosomal regions and (ii) exhibit a stronger dependence on *smc* for alignment (Fig. S5C). To test this model, we fluorescently labeled pairs of DNA loci, at equivalent distances from *ori*, but on opposite arms of the chromosome, using an orthogonal ParB/*parS* system (Badrinarayanan et al., 2015b). A pair of DNA loci were engineered at +200 kb and +3842 kb, *i.e*. within the *parS-*proximal domain showing strongest SMC enrichment and highest inter-arm Hi-C values. Another pair of DNA loci were labeled in the middle of each arm (+1000 kb and +3042 kb). And finally, to investigate chromosomal arm alignment near *ter*, a pair of DNA loci at +1800 kb and +2242 kb were fluorescently labeled. We then measured, for each pair of loci, inter-focus distances in a population of otherwise WT and Δ*smc* cells (Fig. 3E). To allow cells to elongate and expand their chromosomes, we measured inter-focus distances in cells where the only copy of *dnaA* was driven by a vanillate-regulated P_*van*_ promoter. Washing cells free of vanillate produced a population of cells that each contained just one copy of the chromosome and continued to grow but were unable to divide, leading to an elongated cell where the chromosome fills the entire available cytoplasmic space (Le and Laub, 2016).

As cell length increased with time, we observed that the mean inter-focus distances for each pair of DNA loci also increased, consistent with an overall expansion of the chromosome (Fig. 3E). However, the rate of expansion was different, depending on genomic locations of labeled DNA loci. The inter-focus distance for the *parS*-proximal, 200 kb-3842 kb pair increased only modestly, ~0.5 µm as the cell length tripled (Fig. 3E). In contrast, the inter-focus distance for mid-arm loci increased ~1.5 µm on average as the cell length tripled (Fig. 3E). Importantly, the inter-focus distance of the *parS*-proximal loci, but not the other locus pairs, increased in the *Δsmc* background (Fig. 3E). These microscopy-based analyses support our hypothesis that SMC most effectively aligns the chromosome arms nearest the *ori* and *parS* site where it is loaded.

In parallel, we analyzed Hi-C data on the elongated cells resulting from DnaA depletion (Fig. S5D-E) (Le and Laub, 2016). We observed that ~300 kb surrounding *parS* remains well-aligned even in elongated cells 3 hrs after starting the depletion (Fig. S5D-E). In contrast, chromosomal arm alignment elsewhere on the chromosome was rapidly lost to the same extent as in Δ*smc* cells 2 hrs after starting the depletion (Fig. S5F-G). These results agree well with the single-cell microscopy results and together suggest that *Caulobacter* SMC functions most effectively to align chromosomal arms near *ori/parS*.

### Conflicts with transcription influence the SMC-mediated alignment of chromosomal arms

We noted that the region showing the strongest inter-arm interactions did not extend as far from *parS* in the Flip 1-5 strain or in the strains harboring the ectopic *parS* sites near *ter* (Fig. 3C-D, Fig. S5A-B). These observations suggested that the genomic context surrounding *parS* may influence the translocation of SMC along each arm of the chromosome and, consequently, the patterns of inter-arm interaction.

We hypothesized that highly-expressed genes, particularly those oriented toward *parS*, may limit the translocation of SMC. To test this idea, we engineered a strain, hereafter called Flip 2-5, in which a ~419 kb DNA segment between +3611 kb and +4030 kb on the left arm was inverted (Fig. 4A). This inversion leaves *parS* at its original location but dramatically changes the genomic context of the flanking DNA on one side of *parS* while leaving the other side unperturbed (Fig. 4A, Fig. S6A-B). The inverted segment contains several highly-expressed genes, including a rRNA gene cluster and genes encoding ATP synthase, the glycine cleavage system, and cytochrome c oxidase, that all normally read in the *ori-ter* direction, *i.e*. co-directionally with SMC translocation (Fig. 4A, Fig. S6A-B) (Le and Laub, 2016).

To assess the density of RNA polymerases directly, we performed ChIP-seq on exponentially-growing cells producing RpoC*-*FLAG as the only version of RNA polymerase β’ subunit (Fig. 4A, Fig. S6A-B). Separating sequencing reads based on the direction of transcription clearly demonstrated an enrichment of highly-expressed genes transcribed in the *ori-ter* direction in WT cells between +3611 and +4030 kb (Fig. 4A, Fig. S6A). We also confirmed, using ChIP-seq on RpoC-FLAG, that these same genes remain highly expressed in the Flip 2-5 background, but now on the opposite strand such that they read in a *ter-ori* direction (Fig. 4A, Fig. S6B).

The Hi-C contact map of G1-phase Flip 2-5 cells showed, in sharp contrast to WT cells, a pronounced asymmetrical pattern of inter-arm interactions (Fig. 4B, Fig. S7A-B). The inter-arm interactions in the Flip 2-5 strain manifested as a nearly vertical streak in the Hi-C map, still emanating from a *parS*-proximal position (Fig. 4B). This vertical streak indicates that the strongest inter-arm interactions now occur between an ~50-80 kb region of DNA on the left side of *parS* with an ~400 kb segment on the right arm of the chromosome, a pronounced asymmetry compared to the pattern in WT cells (Fig. 4B). We confirmed that this asymmetric pattern of inter-arm interactions in the Flip 2-5 strain is still dependent on SMC as this vertical streak disappeared from the contact map of Flip 2-5 *Δsmc* cells (Fig. 4B). These results suggest that the genomic context of DNA flanking the *parS* site, and likely the orientation of highly expressed genes, can dramatically influence the pattern of inter-arm contacts.

To assess whether the Flip 2-5 strain also led to a change in the genomic distribution of SMC, we performed ChIP-seq on FLAG-SMC in the Flip 2-5 background (Fig. 5, Fig. S4C, Pearson’s correlation coefficient between ChIP-seq and Hi-C data = 0.6, p < 10^-12^). The enrichment of SMC dropped off slightly faster in the first ~150 kb away from *parS*, in the *ori-ter* direction, in the Flip 2-5 strain compared to WT (Fig. 5A and Fig. S7C). However, most strikingly, the Flip 2-5 strain exhibited a series of new SMC ChIP-seq peaks, in addition to the one at *parS*, particularly in the region that was inverted (solid triangles, Fig. 5A-C). Comparing the SMC and RpoC ChIP-seq profiles indicated that these new SMC peaks coincided with the highly expressed gene clusters that had been reoriented in the inversion to read toward *ori* and *parS* (Fig. 5C). These new peaks are not artefacts that are normally associated with highly-transcribed genes since (i) they are unique in Flip 2-5 cells but not in the WT or Flip 1-5 control, and (ii) the shape and the magnitude of the unique peaks in Flip 2-5 are distinct from that of WT and the Flip 1-5 cells (Fig. 5C). These data suggested that transcription in a head-on orientation with SMC translocation can either impede SMC translocation away from *parS*, unlink potential handcuffed SMC, or drive the dissociation of SMC from DNA, limiting the extent of inter-arm contacts and, in some cases, produce an asymmetrical pattern of inter-arm contacts by Hi-C.

To test whether the asymmetrical inter-arm interactions in the Flip 2-5 strain arise because of the reoriented, highly expressed genes within the inverted genomic region, we treated Flip 2-5 cells with rifampicin before fixing cells for Hi-C (Fig. 4B). Rifampicin inhibits transcription elongation in bacteria, thereby eliminating actively translocating RNA polymerases from the chromosome. As we reported previously for WT cells, the inhibition of transcription reduced short-range intra-arm contacts (Le et al., 2013). In addition, for the Flip 2-5 strain, the vertical streak was eliminated and the inter-arm interactions reverted to a symmetric pattern on the diagonal (Fig. 4B, Fig. S7B), demonstrating that transcription is required for an asymmetrical inter-arm interaction pattern.

To further test the relationship between the orientation of highly expressed genes and SMC-dependent inter-arm interactions, we constructed three additional strains with different chromosomal inversions (Fig. 6A). We wondered if reversing the transcription orientation of a single highly-transcribing rRNA gene cluster would be sufficient to induce an asymmetrical inter-arm pattern. To test this possibility, we created the Flip 3-4 strain (Fig. 6A). The Hi-C map of G1-phase cells of the Flip 3-4 strain showed negligible changes to the inter-arm alignment compared to non-flipped cells (Fig. 6B-C). However, the inverted region in this strain is ~240 kb away from *parS* and, as noted above, SMC and SMC-dependent inter-arm interactions are strongest within a limited range around *parS*. Thus, we reasoned that the effect of an inverted rRNA cluster would be stronger if placed closer to *parS*. We did so by constructing the Flip 2-4 strain that has DNA between +3788 kb and +4030 kb inverted (Fig. 6A). Hi-C on G1-phase cells of the Flip 2-4 strain showed a pronounced asymmetrical inter-arm interaction pattern (~20° deviation from a diagonal, starting after 80 kb from *ori*), though less dramatic than that of the Flip 2-5 cells in which several highly expressed genes in addition to the rRNA locus were inverted (Fig. 6B-C, Fig. 4B).

**Figure 6:**
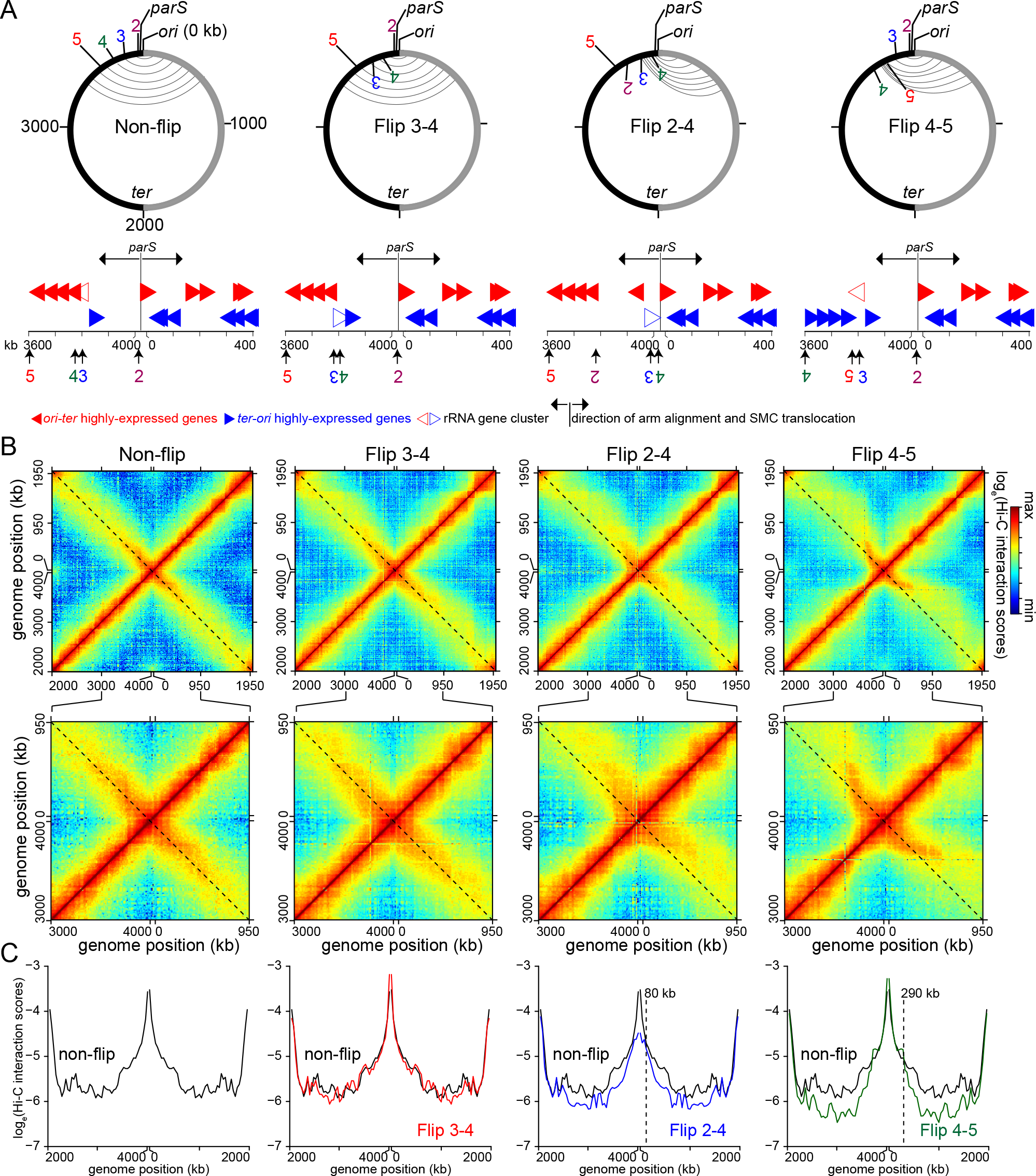
Inverting 180 kb DNA segment containing highly-expressed *ori-ter* genes is sufficient to induce an asymmetrical pattern of inter-arm contacts. **(A)**Schematic genomic maps for WT cells (non-flip) and the inversion strain Flip 3-4, Flip 2-4 and Flip 4-5. The inversion end points (2, 3, 4 and 5) together with the genomic location of *parS* and *ori* are indicated on the map. The aligned DNA regions, as observed by Hi-C (panel B), are presented schematically as grey curved lines connecting the two chromosomal arms. Below each genomic map are the positions of highly-expressed genes that transcribe in the *ori-ter* (solid red arrows) or *ter-ori* (solid blue arrows) direction in each strain. The positions of rRNA gene cluster are indicated with open red or blue arrows. The genomic position of *parS* and the direction of SMC translocation are shown with black arrows. Only 400-kb DNA segments surrounding *ori* are shown. (B) Normalized Hi-C contact maps for non-flip, Flip 3-4, Flip 2-4 and Flip 4-5 cells. The black dashed line indicates the secondary diagonal of the square matrix. A 1000 kb region surrounding *parS/ori* were zoomed in and presented below each whole-genome Hi-C contact map. (C) Hi-C interaction scores along the secondary diagonal for contact maps of non-flip (black), Flip 3-4 (red), Flip 2-4 (blue), and Flip 4-5 cells (dark green). Vertical black dashed lines show position where Hi-C interaction scores start to reduce in the Flip strains compared to the non-flip strain.

We further investigated the effect of transcription orientation bias on chromosomal arm alignment by inverting a DNA segment between +3611 kb and +3788 kb, the Flip 4-5 strain. Although this section does not contain an rRNA gene cluster, it includes four highly-transcribed operons that normally transcribe in the *ori-ter* direction in WT cells (Fig. 6A, denoted with square brackets in Fig. 4A). Preceding this segment, DNA between +3788 kb and +4030 kb is largely free of highly-expressed genes oriented toward *ori/parS* (Fig. 4A). Hi-C on G1-phase cells of the Flip 4-5 strain showed two distinct phases of inter-arm contacts (Fig. 6B-C). The first phase (~290 kb) is a typical set of symmetrical inter-arm contacts as seen in WT cells. The second phase coincides with the inverted DNA segment and has a pronounced asymmetrical inter-arm pattern (Fig. 6B-C). Collectively, our results emphasize that highly-transcribed genes, depending on their transcriptional direction, can dramatically influence the action of SMC and the global organization of a chromosome.

## DISCUSSION

### Conflicts between SMC and RNA polymerase, and the consequences for bacterial chromosome organization

Chromosomes in all organisms are typically laden with DNA-bound proteins that likely influence the dynamics and movement of SMC (Ocampo-Hafalla et al., 2016; Stigler et al., 2016; Wang et al., 2017). Our results here indicate that the distribution and translocation of *Caulobacter* SMC, and the consequent alignment of chromosomal arms, is strongly influenced by actively transcribed genes, particularly those oriented toward *ori*. The observed chromosome organization defect likely stems from a conflict between SMC and transcription, without involving the replisome. This conclusion is based on two key observations. First, the experiments involving strains with inverted chromosomal regions were performed on synchronous G1-phase cells, *i.e*. non-replicating cells. Second, the Flip 1-5 strain, where the SMC loading site was relocated to a mid-arm position, has highly-expressed genes (such as the rRNA gene cluster) transcribing in the opposite direction to that of the replisome in actively replicating cells. However, this strain did not exhibit a dramatic off-diagonal pattern of DNA interactions (Fig. 3C). Thus, we suggest that head-on conflicts between translocating SMC and RNA polymerase can directly shape bacterial chromosome organization.

The mechanism(s) that drive translocation of bacterial SMC from *parS* remains unknown. It is tempting to speculate that the *ori-ter* transcription bias (Fig. S6A) might indicate that transcription helps push SMC away from its *ori*-proximal loading site. However, chromosome arm alignment in WT *Caulobacter* cells treated with rifampicin is not reduced (Fig. S7D-E). *E. coli* MukBEF, a non-canonical SMC, has been proposed to translocate via a “rope climber” mechanism (Badrinarayanan et al., 2012). In this model, a concerted opening and closing of just one MukBEF dimer in a handcuffed dimer-dimer allows an opened dimer to grab the next DNA segment before releasing the previously closed MukBEF, thereby “swinging” the dimer-dimer complex down DNA (Badrinarayanan et al., 2012). In *B. subtilis*, recent structural studies suggested that ATP binding, hydrolysis, and release can switch SMC between a rod- and a ring-like conformational states, with this motor-like cycling somehow mediating SMC translocation (Bürmann et al., 2013; Minnen et al., 2016). However, none of these current models can explain the *directional* movement of SMC from its loading site.

### SMC loading at the bacterial centromere parS site: a coupling between chromosome organization and segregation

Our results support a model in which *Caulobacter* SMC is, like in *B. subtilis*, recruited at or near *parS*, loaded in a ParB-dependent manner, and then redistributed towards *ter*. ParB is a bacterial-specific protein, but likely works closely with and coevolves with SMC to ensure chromosome organization and chromosome segregation in bacteria. Interestingly, *S. pneumoniae* lacks a ParA homolog, yet retains ParB-*parS* to recruit SMC (Minnen et al., 2011), underscoring the tight connection of ParB and SMC. It is worth noting that, unlike *B. subtilis*, deleting *smc* in a wide range of bacteria, including *Caulobacter*, does not cause sensitivity to high temperature or fast-growing conditions (Le et al., 2013; Nolivos and Sherratt, 2014). Conversely however, ParA-ParB-*parS* is essential in *Caulobacter* but not in *B. subtillis* (Mohl and Gober, 1997; Murray and Errington, 2008). The ParA-ParB-*parS* system and the SMC complex likely collaborate to ensure proper chromosome segregation and organization, but with slightly different contributions or relative weights in different organisms. Finally, there is a distant SMC homolog, called the MukBEF system, in γ-proteobacteria (Nolivos and Sherratt, 2014). Notably, ParA-ParB-*parS* systems do not exist in these bacteria, leaving open the question of how MukBEF complex loads on the chromosome, assuming it requires a specific loader (Nolivos and Sherratt, 2014). *E. coli* MukBEF was found, by ChIP-seq, to enrich at *ter* (Nolivos et al., 2016). We also observed an enrichment of *Caulobacter* SMC near the *ter* region (Fig. 3A-B), and a slight increase in inter-focus distance in *Δsmc* cells when intra-arm loci at +1600 kb and +1800 kb were labeled (Fig. 3E). The enrichment of *Caulobacter* SMC near *ter* might occur via a *parS*-independent *E. coli*-like mechanism, however it does not result in DNA alignment in this area (Fig. 1B). The mechanism of SMC enrichment at *ter* is currently unknown in *Caulobacter*.

### Evidence that bacterial SMC tethers chromosomal arms together

The *Caulobacter* SMC complex promotes interactions between loci at approximately equivalent positions on opposite arms of the chromosome up to at least 600 kb from *parS*. It could be that SMC physically tethers the arms together. Alternatively, SMC could promote alignment passively by compacting each arm separately, reducing the cytoplasmic mobility of each arm and thereby increasing inter-arm interactions. The Hi-C patterns documented here for the WT and various inversion strains are most easily explained by the active alignment model in which SMC physically links DNA from both arms together (Fig. 7). Such a model is also appealing given the notion that SMC can topologically entrap DNA. Moreover, contact probability curves derived from the Hi-C data, which reflect global chromosome compaction, were generally very similar for *Δsmc* and WT cells (Le et al., 2013), suggesting that SMC plays only a minor role in intra-arm compaction in *Caulobacter*. Nevertheless, we cannot completely rule out other possible ParB-independent roles of SMC on chromosomal arm compaction. In the active alignment model, also suggested by studies of *B. subtilis* SMC (Wang et al., 2017), the inter-arm interactions documented by Hi-C may reflect loop generation by SMC (Fig. 7). In WT cells, DNA from each chromosomal arm may be effectively threaded through SMC at approximately similar rates as SMC moves towards *ter* and away from *parS* (Fig. 7A-B). In the inversion strains, DNA on the left arm may be threaded through SMC less efficiently than the right arm due to conflicts with convergent transcription (Fig. 7C). As suggested for eukaryotic SMC, loop enlargement may be a general mechanism for folding chromosomes or bringing distant loci together (Fudenberg et al., 2016; Nasmyth, 2001).

**Figure 7:**
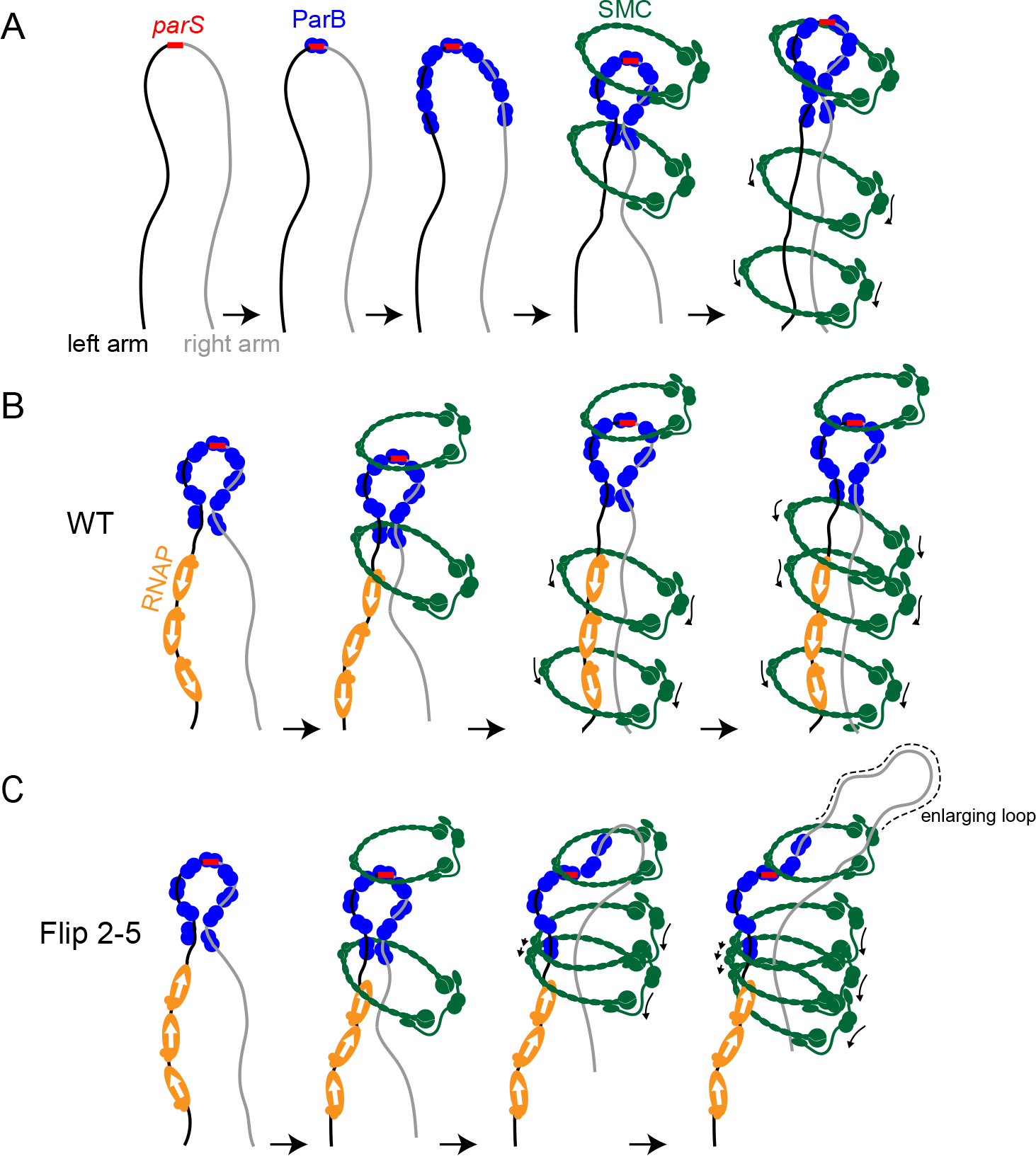
A schematic model for an active alignment of the left and right chromosomal arms by SMC. **(A)** ParB (blue) binds to the bacterial centromere *parS* site (red), spreads and might bridge the left (black) and the right (grey) chromosomal arms together. SMC (dark green) is recruited by ParB and most likely tether the two arms of the chromosome together. An SMC-ScpA-ScpB complex can either hold both chromosome arms within its lumen or two SMC complexes, each encircles one chromosome arm can handcuff to tether both arms together. For simplicity, only SMC encircling both arms are shown schematically. **(B-C)** A schematic model of how a high density of converging RNA polymerases (orange) might interact physically with SMC or create unfavourable DNA supercoiling that stalls or dissociates SMC from the left chromosomal arm as can happen in the Flip 2-5 strain. Schematic pictures are not to scale.

### Caulobacter SMC is not uniformly distributed across the genome

*Caulobacter* SMC may translocate only ~600 kb toward *ter* while *B. subtilis* SMC loaded at *parS* translocates the full length of the chromosome to the terminus. It is possible that *Caulobacter* SMC may suffer more frequent conflict with convergent transcribed genes (see direction bias ratio-Fig. S6A), or be more sensitive to dissociation following such conflicts, leading to less extensive arm-arm interaction. The extent to which the two arms "zip up" may not matter. If the primary role of SMC-mediated arm-arm interactions is to help enforce the individualization of sister chromosomes immediately after DNA replication, it may only be necessary to ensure that SMC can cohese *parS*-proximal regions of each chromosome. Indeed, in *Caulobacter*, ParA-ParB-*parS* are only required for the segregation of *ori*-proximal DNA, but not of the distal DNA loci (Badrinarayanan et al., 2015b). Once the *ori*-proximal DNA is properly segregated, by SMC and ParA-ParB-*parS*, distal DNA regions follow suit, driven by separate molecular machinery, or more likely without the need of a dedicated system (Badrinarayanan et al., 2015b). In such a case, it may be sufficient to have SMC tether together only a limited region of DNA flanking the *parS* sites. This model would imply that conflicts between SMC and highly expressed genes oriented toward *ori* are most detrimental, with respect to chromosome segregation, if they occur in close proximity to *parS*. Testing this model and further understanding the relationship of SMC and gene expression and its influence on chromosome organization is an important challenge for the future.

## EXPERIMENTAL PROCEDURES

For all experimental procedures, see the Supplementary Information

## AUTHOR CONTRIBUTIONS

Conceptualization, N.T.T., M.T.L., and T.B.K.L.; Data analysis, N.T.T., M.T.L., and T.B.K.L.; Writing, N.T.T., M.T.L., and T.B.K.L.; and Funding Acquisition, T.B.K.L.

## ACKNOWLEDGMENTS

We thank Anjana Badrinarayanan, Hugo Brandao, Matt Bush, and Monica Guo for discussion and comments on the manuscript. We thank Lucy Shapiro, Martin Thanbichler, and Susan Schlimpert for materials. This study was supported by a Royal Society University Research Fellowship (UF140053 and RG150448), a BBSRC grant (BB/P018165/1) to T.B.K.L.M.T.L. is an Investigator of the Howard Hughes Medical Institute.

## ACCESSION NUMBERS

The accession number for the sequencing data reported in this paper is GEO: **GSE97330**.

